# Ablation of proliferating neural stem cells during early life is sufficient to reduce adult hippocampal neurogenesis

**DOI:** 10.1101/308387

**Authors:** Mary Youssef, Varsha S. Krish, Greer S. Kirshenbaum, Piray Atsak, Tamara J. Lass, Sophie R. Lieberman, E. David Leonardo, Alex Dranovsky

**Author notes:** Address correspondence to AD or EDL, s.

## Abstract

Environmental exposures during early life, but not during adolescence or adulthood, lead to persistent reductions in neurogenesis in the adult hippocampal dentate gyrus (DG). The mechanisms by which early life exposures lead to long-term deficits in neurogenesis remain unclear. Here, we investigated whether targeted ablation of dividing neural stem cells during early life is sufficient to produce long-term decreases in DG neurogenesis. Having previously found that the stem cell lineage is resistant to long-term effects of transient ablation of dividing stem cells during adolescence or adulthood (Kirshenbaum et al., 2014), we used a similar pharmacogenetic approach to target dividing neural stem cells for elimination during early life periods sensitive to environmental insults. We then assessed the Nestin stem cell lineage in adulthood. We found that the adult neural stem cell reservoir was depleted following ablation during the first postnatal week, when stem cells were highly proliferative, but not during the third postnatal week, when stem cells were more quiescent. Remarkably, ablating proliferating stem cells during either the first or third postnatal week led to reduced adult neurogenesis out of proportion to the changes in the stem cell pool, indicating a disruption of the stem cell function or niche following stem cell ablation in early life. These results highlight the first three postnatal weeks as a series of sensitive periods during which elimination of dividing stem cells leads to lasting alterations in adult DG neurogenesis and stem cell function. These findings contribute to our understanding of the relationship between DG development and adult neurogenesis, as well as suggest a possible mechanism by which early life experiences may lead to lasting deficits in adult hippocampal neurogenesis.

## Introduction

Adult hippocampal neurogenesis, which occurs in the dentate gyrus (DG), has been the topic of significant study to understand its regulation and function in health and disease (Cameron and Glover, 2015; Ming and Song, 2005). While the rodent DG begins to form during the late embryonic period, most of the structure develops during the first two postnatal weeks (Altman and Bayer, 1990; Angevine, 1965; Bayer, 1980; Stanfield and Cowan, 1979). The majority of granule neurons that comprise the adult rodent DG granule cell layer (GCL) are born during the first week of life (Angevine, 1965; Muramatsu et al., 2007). Accordingly, DG cell proliferation is dramatically higher during this time window compared to later developmental periods (Bayer and Altman, 1974; Muramatsu et al., 2007; Navarro-Quiroga et al., 2006). Throughout the lifelong DG development, new granule neurons arise from Nestin and glial fibrillary acidic protein (GFAP) expressing stem astrocytes (Zhao et al., 2008) and are added to the GCL. During the first postnatal week, these neural stem cells are diffusely spread in the area that will become the GCL, but as the structure matures, the stem cells condense into the subgranular zone (SGZ) where they remain into adulthood (Nicola et al., 2015; Sugiyama et al., 2013). The DG develops in an outside-in manner, with embryonic- and very early postnatal-born cells residing in the outer layer of the GCL and later-born cells filling in the middle and inner layers; adult born cells reside in the inner layer adjacent to the SGZ (Mathews et al., 2010; Muramatsu et al., 2007).

Adult neurogenesis is likely a continuation of the developmental process in the DG (Encinas et al., 2013; Kuhn and Blomgren, 2011; Nicola et al., 2015). By the third week of life in rodents, levels of DG cell proliferation are closer to adult levels than to early developmental levels (Bayer and Altman, 1974; Stanfield and Cowan, 1979). Granule cells born on postnatal day (P) 14 or later are mostly restricted to the inner layer of the GCL, where adult born cells are found (Stanfield and Cowan, 1979). Further, by P14 DG stem cells are condensed into the SGZ where adult stem cells reside (Nicola et al., 2015; Sugiyama et al., 2013). Thus, some studies suggest that “adult” neurogenesis may begin as early as P14, when the DG more closely resembles the adult structure in terms of neurogenesis and anatomical layout (Nicola et al., 2015). Ongoing changes in DG neurogenesis and structure across its life-long development may underlie the short and long-term effects of environment on this structure.

Environmental exposures during early life can alter DG development, often resulting in persistent deficits in DG neurogenesis and function. Early life insults can be caused by medicinally-utilized drugs, illicit substances, infections and psychological stress, among other things. Administering anesthetics or glucocorticoids during the first three postnatal weeks leads to decreased DG cell proliferation and neurogenesis, depletion of the stem cell pool, and increased apoptosis in rodent models (Lu et al., 2017; Yan et al., 2017; Yu et al., 2010; Zhu et al., 2010). Moreover, the acute reductions in DG cell proliferation and stem cell number caused by early postnatal isoflurane and dexamethasone administration persist for several weeks after the exposure and even into adulthood (Yu et al., 2010; Yu et al., 2017b; Zhu et al., 2010). Similarly, ethanol administration and measles infection result in reductions in DG neural stem/precursor cells and neurogenesis, with concurrent increases in apoptosis shortly after the interventions (Fantetti et al., 2016; Xu et al., 2015). Further, early life stress caused by maternal separation or erratic maternal care results in decreased DG cell proliferation and neurogenesis shortly after the stress (Baek et al., 2011; Baek et al., 2012; Lajud et al., 2012; Oreland et al., 2010), with the deficits continuing into adulthood (Aisa et al., 2009; Hulshof et al., 2011; Kikusui and Mori, 2009; Leslie et al., 2011; Mirescu et al., 2004; Naninck et al., 2015; Suri et al., 2013). The young hippocampus is particularly susceptible to many of these harmful interventions, as similar insults during adolescence or adulthood do not produce as severe or lasting effects on DG stem cells or neurogenesis (Elizalde et al., 2010; Heine et al., 2004; Lagace et al., 2010; Zhu et al., 2010). Together, the data suggest that environmental exposures that acutely alter DG development during early life result in long-term and extensive effects on DG neurogenesis.

The direct contribution of the immediate depletion of stem cells and reduction in cell proliferation and neurogenesis to the long-term effects on DG neurogenesis caused by these early life interventions remains unclear. Recent work showed that ablation of dividing stem cells, and consequently reduction of cell proliferation and neurogenesis, for one week during adolescence or adulthood does not have a lasting effect on the DG stem cell lineage (Kirshenbaum et al., 2014). These results parallel studies of drug administration or stress during similar time periods in which there is a recovery of neurogenesis after termination of the intervention (Elizalde et al., 2010; Heine et al., 2004; Lagace et al., 2010; Zhu et al., 2010). Given the prolonged effects of early life interventions, we hypothesized that the immediate reductions in stem cell number, cell proliferation and neurogenesis caused by early postnatal insults contribute to the resulting longterm deficits in DG stem cells and neurogenesis since these changes are induced during critical DG developmental periods. Although studies have shown that ablation of proliferating stem cells during or after the fifth postnatal week does not have lasting effects on DG stem cell lineages (Kirshenbaum et al., 2014), it is unknown if a similar intervention during earlier periods can lead to long-term changes. Here, we examined whether targeting dividing neural stem cells for elimination for one week during early life is sufficient to diminish adult neurogenesis, and if the timing of the intervention affects the outcome. We focused on the first and third postnatal weeks because studies of early life interventions commonly used in rodents encompass one or both of these periods and the development and structure of the DG differs during these two time windows in terms of cell proliferation, birth of surviving neurons, and anatomical structure.

We explored the effect of ablating neural stem cells dividing during the first and third postnatal weeks, then assessed adult DG stem cell lineages and neurogenesis. We found that pharmacogenetic ablation of proliferating GFAP+ cells during the first postnatal week led to fewer adult stem cells and nearly absent neurogenesis, while ablation of proliferating GFAP+ cells during the third postnatal week resulted in reduced neurogenesis, but did not deplete the stem cell pool. Measuring baseline stem cell proliferation during these two time windows revealed that there are more proliferating stem cells during the first postnatal week compared to the third week. Thus, fewer adult stem cells present following the first week intervention may reflect depletion of the stem cell pool. Similar effects of our pharmacogenetic intervention were identified in male and female mice.

## Materials and Methods

### Animals

All mice were maintained on a 12:12 light cycle (6 am to 6 pm light) and given food and water *ad libitum*. All procedures were performed in the light cycle and complied with NIH AALAC and NY State Psychiatric Institute IACUC guidelines.

#### Breeding

A triple-gene modified mouse line was used for this study. Transgenic mice expressing the gene for herpes thymidine kinase under control of the GFAP promoter (GFAP-Tk; RRID:IMSR_JAX:005698) (Bush et al., 1998) were crossed to transgenic mice expressing the gene for Cre-ER^T2^ under control of the Nestin promoter (Nestin-CreER^T2^) (Dranovsky et al., 2011). GFAP-Tk and Nestin-CreER^T2^ animals were maintained on a Cre reporter line homozygous for enhanced yellow fluorescent protein gene preceded by a floxed stop sequence knocked into the ROSA26 locus (R26R-stop-EYFP) (Srinivas et al., 2001). GFAP-Tk^+/-^, Nestin-Cre^+/-^, R26R-EYFP^+/+^ female mice were crossed to GFAP-Tk^-/-^, Nestin-Cre^-/-^, R26R-EYFP^+/+^ male mice to produce littermate GFAP-Tk^-/-^ and ^+/-^, Nestin-Cre^-/-^ and ^+/-^, R26R-EYFP^+/+^ male and female experimental animals. Mice were maintained on a mixed C57Bl/6;Sv129 background. Genotypes were established using PCR as described previously (Bush et al., 1998; Dranovsky et al., 2011; Srinivas et al., 2001).

#### VGCV Treatment

GFAP-Tk^+/-^ experimental mice and their littermate Tk^-/-^ controls were treated with valganciclovir (VGCV, Aurobindo Pharma) from P0-P7 or from P14-P21. For the P0-P7 group, pups were cross-fostered from GFAP-Tk dams to FVB/NJ (Jackson Laboratory, Cat# 001800) foster dams between 4-6 pm on P0 (defined as the day pups were found). FVB/NJ dams were fed chow containing 700 mg VGCV/kg chow (Custom Animal Diets) starting at least 3 days prior to cross-foster. VGCV chow was available *ad libitum* to FVB/NJ foster dams until the evening of P7, when it was replaced with standard chow. FVB/NJ dams consumed an average of 10 g of chow on P0, steadily increasing to 14 g chow on P7, thereby receiving 7-10 mg of VGCV per day across the week. Pups were weaned on P28. For the P14-P21 VGCV treatment group, pups were kept with their original mothers. Pups underwent oral gavage with VGCV (10 mg/kg per dose suspended in dH2O) twice daily 8 hours apart starting from the evening of P14 until the evening of P21. Pups were weaned on P28. Mice were separated by sex at weaning. Tk- and Tk+ animals were cohoused in the same cages.

#### CldU Injections, Tamoxifen Administration, and Euthanasia

For the P0-P7 VGCV group, chlorodeoxyuridine (CldU, Sigma-Aldrich, Cat# C6891) was injected intraperitoneally (i.p.) at a dose of 42.5 mg/kg once daily on P6 and P7 between 4-6 pm. For the P14-P21 vGCV group, CldU (42.5 mg/kg) was injected i.p. once daily on P20 and P21 between 4-6 pm.

Tamoxifen (TMX, Sigma-Aldrich, Cat# T5648) was suspended in 1:1 honey:water solution and administered by oral gavage to Nestin-Cre^+/-^ experimental animals once daily on P56 and P57 at a dose of 2.5 mg/animal per day.

For the immediate post-treatment experiments, animals were euthanized on the last day of VGCV treatment (P7 or P21). For the one week recovery experiments, animals were euthanized one week after completion of VGCV treatment (P14 or P28). For the lineage analysis studies, animals were euthanized 7 weeks after TMX administration (P104).

#### Nestin-Kusabira Orange Mice

Transgenic mice expressing a PEST-tagged Kusabira Orange fluorescent protein under control of the Nestin promoter (Nestin-KOr) (Kanki et al., 2010) were used to visualize stem cells. Mice were maintained on a C57Bl/6 background. Nestin-KOr^+/+^ males were crossed to C57BL/6J females (Jackson Laboratories, Cat# 000664) to generate Nestin-KOr^+/-^ experimental animals. Experimental mice were euthanized on P7 or P21. Genotypes were established using PCR as described previously (Kanki et al., 2010).

### Tissue Preparation

Mice were anesthetized with a mixture of 150 mg/kg ketamine and 10 mg/kg xylazine and perfused transcardially with ice cold phosphate buffered saline (PBS), followed by 4% paraformaldehyde (PFA) in PBS. Brains were removed and postfixed by immersion in 4% PFA for 24 hours, then switched to 30% sucrose in PBS for 48 hours for cryoprotection. Whole heads for animals euthanized at ages P7, P10, P14, P21 and P28 were stored in 4% PFA for an additional 24 hours before brain extraction to minimize damage during removal from skull. Brains were sectioned coronally on a cryostat to prepare 35 μm sections, which were stored in PBS with 0.02% sodium azide at 4°C. Tissue was serially sectioned into six wells per brain.

### Immunohistochemistry

Free floating tissue sections were triple washed in PBS, pre-treated with 2N hydrochloric acid/0.1M boric acid or 1% triton and triple washed in PBS again if necessary (see below), then incubated for 1 hour in blocking solution (12.5% normal donkey serum and 0.25% triton in PBS), followed by incubation with one or more of the following primary antibodies diluted in blocking solution: chicken anti-GFP 1:500 (Abcam, Cat# ab13970, RRiD: AB_300798), goat anti-DCX 1:250 (Santa Cruz Biotechnology, Cat# sc-8066, RRID: AB_2088494), goat anti-GFAP 1:500 (Abcam, Cat# ab53554, RRID: AB_880202), goat anti-MCM2 1:100 (Santa Cruz Biotechnology, Cat# sc-9839, AB_648841), mouse anti-GFAP 1:500 (Millipore, Cat# MAB3402, RRID: AB_94844), mouse anti-Nestin 1:100 (BD Biosciences, Cat# 556309, RRID: AB_396354), mouse anti-NeuN 1:1000 (Millipore, Cat# MAB377, RRID: AB_2298772), rabbit anti-GFAP 1:500 (Dako, Cat# Z033429-2, RRID: AB_10013382), rabbit anti-Ki67 1:500 (Abcam, Cat#: ab15580, RRID: AB_443209), rabbit anti-Kusabira Orange 1:500 (MBL International, Cat# PM051, RRID: AB_10597258), rabbit anti-S100β 1:1000 (Abcam, Cat# ab41548, RRID: AB_956280), rat anti-BrdU 1:2500 (Accurate Chemical, Cat# OBT0030G, RRID: AB_2313756). Tissue was incubated with most primary antibodies overnight at 4°C. Stains including anti-BrdU antibody were incubated for 40 hours at 4°C. Tissue was incubated with anti-Nestin antibody for seven days at 4°C, and the other antibodies in the stain were added on the sixth day. Following primary antibody incubation, tissue was triple washed in PBS and incubated for one hour with fluorescent secondary antibodies (Jackson ImmunoResearch) diluted 1:400 in PBS with Hoechst 1:10,000 (ThermoFischer Scientific, Cat# 33342). After secondary antibody incubation, tissue was triple washed in PBS, mounted onto slides and coverslipped with Aqua-Poly/Mount (VWR, Cat# 87001-902).

Tissue being stained with anti-BrdU antibody underwent antigen retrieval by pre-treatment with 2N hydrochloric acid (HCl) at 37°C for 30 minutes, followed by 0.1M boric acid at room temperature for 10 minutes. Tissue being stained with anti-NeuN antibody was pre-treated with 1% triton for 30 minutes at room temperature. For more detailed information about staining protocols and antibodies used, see Tables S1-S3.

### Imaging and Cell Quantification

#### GFAP-Tk One-week Recovery Mice

Three (for CldU counts) and five (for Ki67 counts) sections per animal atlas matched (Allen Mouse Brain Atlas) and equally spaced across the anterior-posterior axis of the DG were selected by visualizing Hoechst counterstain using a 10x objective on an epifluorescence microscope (Olympus IX71). CldU+ and Ki67+ cells were manually counted during live visualization of brain slices using a 40x objective on the epifluorescence microscope. CldU+ cells were counted in the entire SGZ and GCL of chosen sections. Ki67+ cells were counted in the entire SGZ of chosen sections. Number of cells was averaged across sections and reported as average number of cells per section.

#### GFAP-Tk Adult Mice

Three sections per animal atlas matched and equally spaced across the anterior-posterior axis of the DG were selected with a 10x objective on the epifluorescence microscope. The entire DG of each chosen section was imaged using the 40x objective on a confocal microscope (Leica TCS SP8). Cells were quantified from images visualized on Leica LAS X Core confocal software (http://www.leica-microsystems.com/products/microscope-software/details/product/leica-las-x-Js/). All EYFP+ cells or all GFAP+EYFP+ cells in chosen sections were manually counted throughout the SGZ, GCL, molecular layer and hilus, and marker co-expression and morphology were manually determined for each cell. EYFP expression facilitated cell quantification as EYFP labels the nucleus, cell body, and processes. Data are reported as sum of cells counted per animal.

#### Nestin-KOr Mice

Three sections per animal atlas matched and equally spaced across the anterior-posterior axis of the DG were selected using Hoechst counterstain with the 10x objective on the epifluorescence microscope. For P7 animals, one 63x frame was captured using the confocal microscope for each blade (upper and lower) of the DG for each section for a total of six frames. For P21 animals, three 63x frames were captured for each blade of the DG for each section. Confocal images were visualized using Leica LAS X Core software. All KOr+GFAP+ cells were manually counted in all captured frames and the number of these cells also expressing MCM2 was manually counted. Two to three hundred KOr+GFAP+ cells were counted per animal.

### Statistical Analysis

Data were analyzed using GraphPad Prism Software (https://www.graphpad.com/scientific-software/prism/, RRID: SCR_002798). All data sets are presented as mean ± standard error of the mean (SEM). Statistical significance was set at p≤0.05.

Immunofluorescence data were analyzed using two-way ANOVA with two fixed factors: condition (genotype: Tk-vs. Tk+; sacrifice age: P7 vs. P21) and sex (male vs. female) (Table S4). For experiments with VGCV treatment, P0-P7 treated animals were compared to each other and P14-P21 treated animals were compared to each other. Wherever main effect of sex was observed, male and female animals were analyzed separately using two-tailed, unpaired t-tests to determine the effect of condition within sex. For analyses in which there was no main of effect of sex, the two-way ANOVA main effect of condition was used to compare conditions.

## Results

### Short-term effects of VGCV treatment on cell proliferation

The goal of this study was to determine if temporally restricted ablation of dividing neural stem cells during early life is sufficient to produce long-term effects on adult stem cell lineages in the DG. We utilized a GFAP-Tk (Bush et al., 1998) mouse line crossed to Nestin-CreER^T2^ (Dranovsky et al., 2011) and R26R-stop-EYFP (Srinivas et al., 2001) lines as described previously (Kirshenbaum et al., 2014). Combining the two genetic systems has allowed us to assess the adult stem cell lineage in animals that have undergone prior stem cell ablation (Kirshenbaum et al., 2014).

Treatment of GFAP-Tk mice with valganciclovir (VGCV) decreases cell proliferation and neurogenesis by targeting dividing stem cells for apoptosis (Garcia et al., 2004). We first sought to determine if our VGCV treatment is effective in reducing cell proliferation and if cell proliferation recovers after cessation of treatment during our experimental time periods as with treatment later in life (Kirshenbaum et al., 2014). We administered VGCV to pups for one week from P0-P7 or from P14-P21 (Fig. 1A, F). VGCV requires oral administration, which created technical challenges for treating nursing pups. For the P0-P7 VGCV treatment, dams were given chow containing VGCV allowing drug delivery via milk to the pups. Since GFAP-Tk+ males are infertile (Bush et al., 1998), pups were cross-fostered to non-transgenic FVBN/J nursing dams in order to avoid the effects of VGCV in Tk+ dams. For P14-P21 VGCV treatment, pups were gavaged twice daily with VGCV to ensure proper dosing because nursing declines and ceases during this time period. Short-term effects of VGCV treatment were established by assessing the number of cells that were dividing at the end of treatment and those dividing one week later. To this end, chlorodeoxyuridine (CldU) was administered intraperitoneally on the last two days of VGCV treatment and animals were sacrificed one week later (Fig. 1A, F). CldU labeling was assessed to identify cells dividing at the end of treatment. The mitotic antigen Ki67 was used to quantify cell proliferation one week after treatment cessation. Despite the potential for variability in amount of drug delivered to pups especially with delivery via milk, there was little variance seen within groups for the short-term effects of VGCV. Male and female animals were analyzed separately because two-way ANOVA revealed a main effect of sex in one of the primary outcome measures (Table S4).

**Figure 1.**
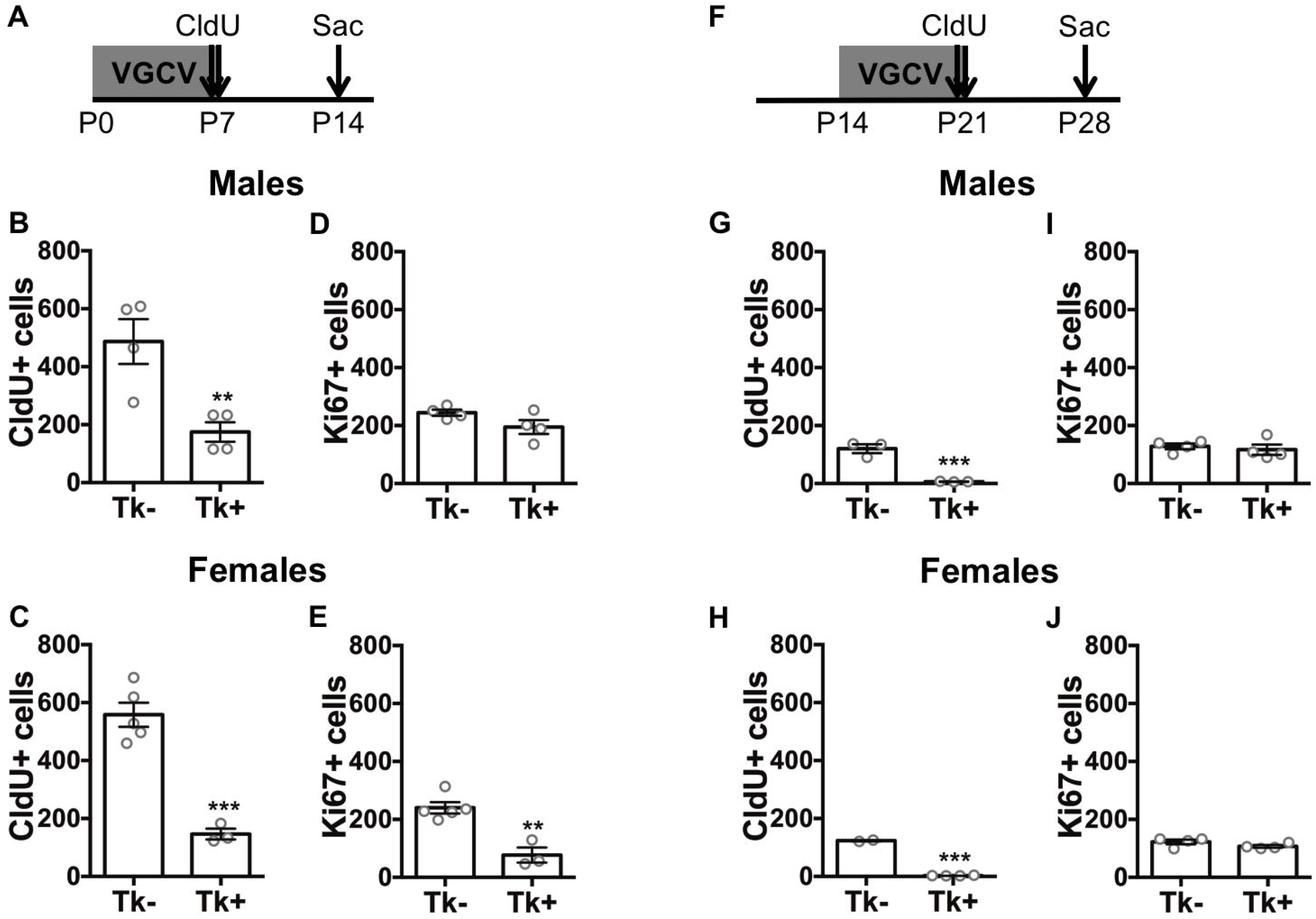
Reduction of cell proliferation in the DG of GFAP-Tk mice by VGCV. (A) Experimental timeline of P0-P7 valganciclovir (VGCV) treatment and sacrifice one week after treatment completion in GFAP-Tk+ animals and Tk-controls. (B) The number of CldU+ cells was reduced in Tk+ males and (C) Tk+ females compared to Tk-controls. (D) No difference in the number of Ki67+ cells was detected between Tk- and Tk+ males. (E) The number of Ki67+ cells was reduced in Tk+ versus Tk-females. (F) Experimental timeline of P14-P21 VGCV treatment and sacrifice one week after treatment completion. (G) The number of CldU+ cells was reduced in Tk+ males and (H) females compared to Tk-animals. (I) No difference in the number of Ki67+ cells was detected between Tk- and Tk+ male or (J) female animals. Data are expressed as mean ± SEM. **p<0.01, ***p<0.001

**Figure 2.**
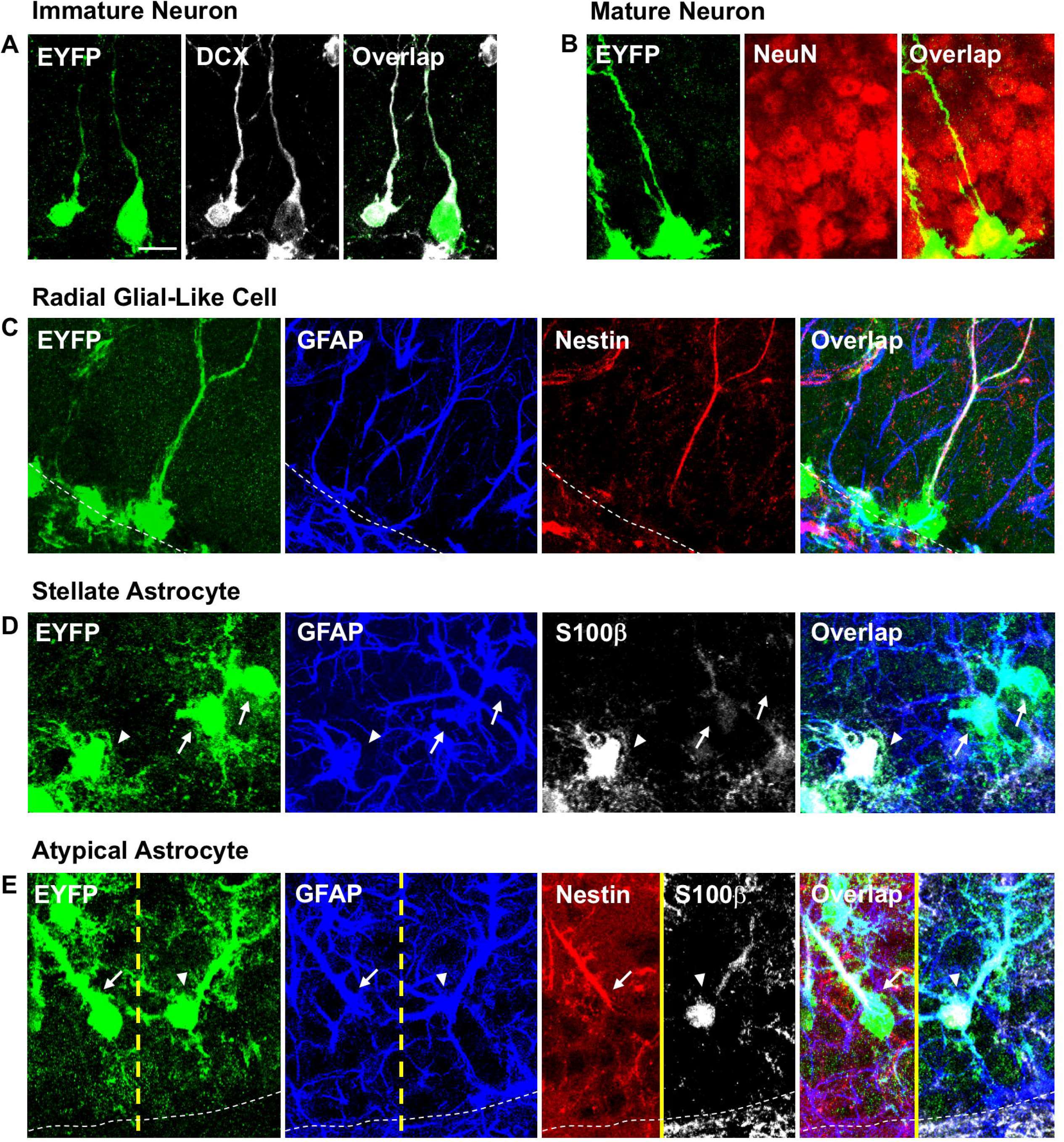
Marker expression and morphology of neurons and astrocytes in the adult Nestin lineage. (A) Representative images of immature neurons co-expressing EYFP and doublecortin (DCX). (B) Representative images of a mature neuron co-expressing EYFP and NeuN. (C) Representative images of a radial glial-like cell (RGL) co-expressing EYFP, GFAP, and Nestin with cell body in the subgranular zone (SGZ) and radial process traversing the granule cell layer. (D) Representative images of stellate astrocytes (SA) co-expressing EYFP and GFAP with multiple similar-sized processes extending from the cell body. Arrowhead points to S100β+ SA. Arrows point to S100β-SAs. (E) Representative images of atypical astrocytes (AA) co-expressing EYFP and GFAP with a single dominant process extending from the cell body, which are in the outer GCL. Arrow points to Nestin+ AA with one process. Arrowhead points to S100β+ AA with one dominant process and multiple minor processes. Scale bar represents 10μM. Dotted lines indicate SGZ.

VGCV treatment led to a robust reduction in the number of CldU+ cells in the SGZ and GCL in Tk+ males (t_(6)_ = 3.709, p=0.01) and females (t_(6)_ = 7.221, p=0.0004) treated from P0-P7 compared to Tk-animals (Fig. 1B, C). The number of Ki67+ cells in the SGZ was used to determine if cell proliferation recovers one week after cessation of VGCV treatment. There was no difference detected in the number of Ki67+ cells between Tk- and Tk+ males treated with VGCV from P0-P7 (t_(6)_ = 1.865, p=0.1114) (Fig. 1D). However, Tk+ females had fewer Ki67+ cells compared to Tk-females (t_(6)_ = 5.062, p=0.0023) (Fig. 1E), indicating that in females cell proliferation did not recover one week after treatment cessation, unlike in males. VGCV treatment from P14-P21 led to almost complete elimination of CldU+ cells in both Tk+ males (t_(4)_ = 7.474, p=0.0017) and females (t_(4)_ = 81.39, p<0.0001) compared to Tk-animals (Fig. 1G, H). Moreover, cell proliferation was restored to near baseline levels one week after end of treatment, as no differences were detected in the number of Ki67+ cells between Tk-and Tk+ animals in both P14-P21 VGCV treated males (t_(6)_ = 0.5672, p=0.5912) and females (t_(6)_ = 1.662, p=0.1476) (Fig. 1I, J). These results indicate that one week of VGCV administration from P0-P7 or from P14-P21 effectively reduces cell proliferation in both males and females, and that cell proliferation returns to near normal levels one week after VGCV termination in all groups except P0-P7 treated females.

### Long-term effects of VGCV treatment on the Nestin lineage

After demonstrating that we are able to effectively and, for the most part, transiently decrease cell proliferation in young animals using the GFAP-Tk system, we sought to determine if, unlike later in life (Kirshenbaum et al., 2014), ablating dividing stem cells during early life results in lasting effects on adult neurogenesis and the stem cell lineage. We treated mice with VGCV from P0-P7 or from P14-P21. Subsequently, we administered tamoxifen (TMX) at P56 and P57 to induce Cre-mediated recombination and EYFP reporter expression in Nestin expressing stem and progenitor cells to begin labeling the adult lineage. Animals were sacrificed 7 weeks after TMX administration to allow labeled cells within the lineage to accumulate (Fig. 3A, Fig. 4A).

**Figure 3.**
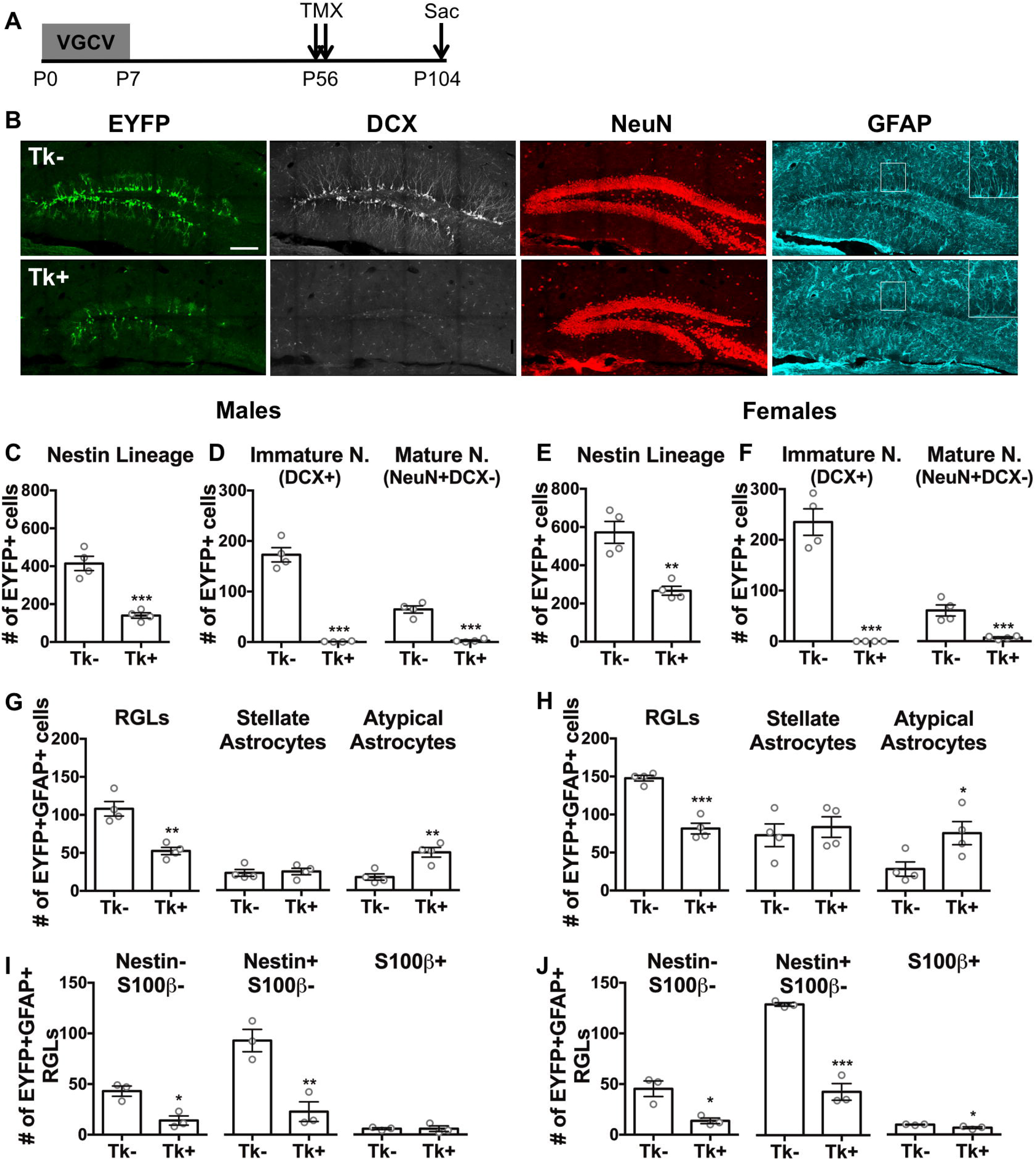
Targeting dividing stem cells from P0-P7 leads to depletion of the DG stem cell pool and decreased neurogenesis in adulthood. (A) Experimental timeline of P0-P7 VGCV treatment and tamoxifen (TMX) administration in GFAP-Tk/Nestin-CreER^T2^ mice. (B) Representative images of EYFP, DCX, NeuN, and GFAP staining in P0-P7 VGCV treated Tk-and Tk+ animals. (C) P0-P7 VGCV led to fewer EYFP+ cells in Tk+ versus Tk-males. (D) P0-P7 VGCV reduced the number of DCX+ immature and NeuN+DCX-mature neurons within the Nestin lineage of Tk+ compared to Tk-males. (E) P0-P7 VGCV led to fewer EYFP+ cells in Tk+ versus Tk-females. (F) P0-P7 VGCV reduced the number of DCX+ immature and NeuN+DCX-mature neurons within the Nestin lineage of Tk+ compared to Tk-females. (G) P0-P7 VGCV decreased the number of RGLs and increased the number of atypical astrocytes, but did not affect the number of stellate astrocytes within the Nestin lineage of Tk+ males and (H) females compared to Tk-animals. (I) P0-P7 VGCV led to fewer Nestin-S100β-and Nestin+S100β-RGLs, but did not change the number of S100β+ (including both Nestin-S100β+ and Nestin+S100β+) RGLs within the Nestin lineage of Tk+ versus Tk-males. (J) P0-P7 VGCV led to fewer Nestin-S100β-, Nestin+S100β-, and S100β+ RGLs within the Nestin lineage of Tk+ versus Tk-females. Scale bar represents 150 μM. GFAP inset is at 2x magnification. Data are expressed as mean ± SEM. *p<0.05, **p<0.01, ***p<0.001

**Figure 4.**
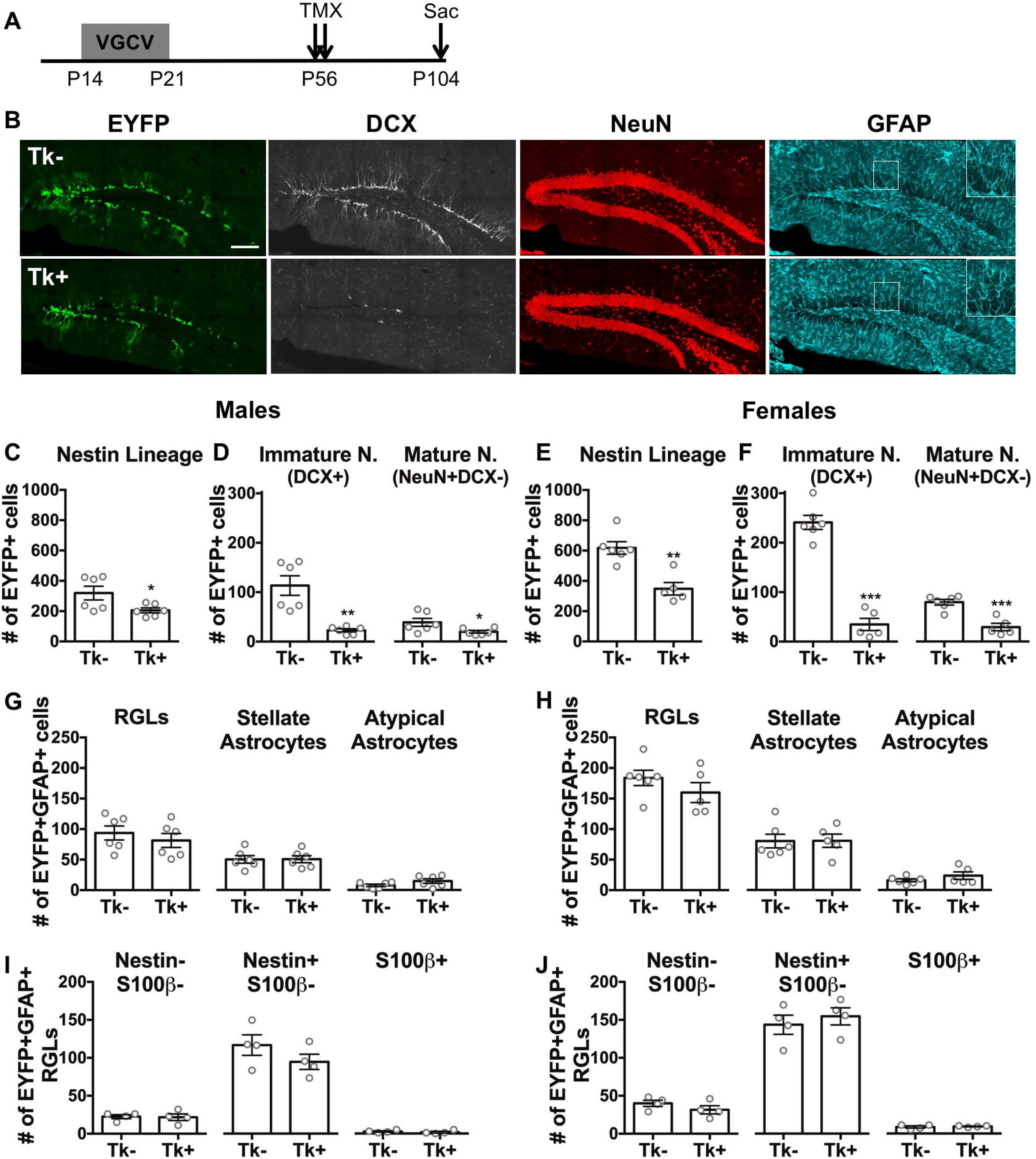
Targeting dividing stem cells from P14-P21 leads to decreased DG neurogenesis, but does not deplete the stem cell pool in adulthood. (A) Experimental timeline of P14-P21 VGCV treatment and TMX administration in GFAP-Tk/Nestin-CreER^T2^ mice. (B) Representative images of EYFP, DCX, NeuN, and GFAP staining in P14-P21 VGCV treated Tk- and Tk+ animals. (C) P14-P21 VGCV led to fewer EYFP+ cells in Tk+ versus Tk-males. (D) P14-P21 VGCV reduced the number of DCX+ immature and NeuN+DCX-mature neurons within the Nestin lineage of Tk+ compared to Tk-males. (E) P14-P21 VGCV led to fewer EYFP+cells in Tk+ versus Tk-females. (F) P14-P21 VGCV reduced the number of DCX+ immature and NeuN+DCX-mature neurons within the Nestin lineage of Tk+ compared to Tk-females. (G) P14-P21 VGCV did not affect the number of RGLs, stellate astrocytes, or atypical astrocytes in the lineage of Tk+ males and (H) females compared to Tk-animals. (I) P14-P21 VGCV did not change the number of Nestin-S100β-, Nestin+S100β- or S100β+ RGLs in the Nestin lineage of Tk+ males and (J) females compared to Tk-animals. Scale bar represents 150 μM. GFAP inset is at 2x magnification. Data are expressed as mean ± SEM. *p≤0.05, **p<0.01, ***p<0.001

We assessed the number and identity of cells in the Nestin lineage by quantifying and categorizing EYFP+ cells throughout the DG subregions, including SGZ, GCL, molecular layer and hilus, according to well-established criteria for classifying cells within the adult Nestin DG lineage (Ming and Song, 2005; Zhao et al., 2008) (Fig. S1). Cellular identity was determined based on marker co-expression and morphological characteristics as described previously (Bonaguidi et al., 2011; Dranovsky et al., 2011; Lagace et al., 2007) (Fig. 2). To identify the neuronal and astrocytic components of the lineage, we stained for doublecortin (DCX), NeuN, and GFAP. DCX and NeuN expression was used to identify immature and mature neurons. Since DCX is mostly expressed during the first three weeks after neuronal birth (Brown et al., 2003), we categorized all EYFP+ cells that co-expressed DCX as immature neurons (Fig. 2A). EYFP+ cells that co-expressed NeuN, but not dCx, were considered mature neurons (Fig. 2B). Further, EYFP+ cells that co-expressed GFAP were identified as astrocytes. Since GFAP is expressed by both stem and non-stem astrocytes in the DG, GFAP+ astrocyte types can be distinguished based on their morphology and expression of other markers (Dranovsky et al., 2011; Patrylo and Nowakowski, 1994; Seri et al., 2004). GFAP+ cells with somas in the SGZ and a single radial process traversing the GCL were categorized as radial-glial like cells (RGLs) (Fig. 2C). Within the RGL population, those cells co-expressing the stem marker Nestin are neural stem cells (Bonaguidi et al., 2011; Lagace et al., 2007; Seri et al., 2004). The stem cell potential of Nestin-RGLs remains uncertain (Encinas et al., 2013). EYFP+GFAP+ cells found throughout the DG with multiple similar-sized processes extending from the cell body were classified as non-stem stellate astrocytes (Patrylo and Nowakowski, 1994) (Fig. 2D). Some of these cells also co-expressed the astrocyte marker S100β. A small number of EYFP+GFAP+ cells did not exhibit the typical RGL or stellate astrocyte morphology. These cells had a single dominant process and occasionally minor processes extending from the cell body, the somas were found outside the SGZ, and the dominant process was often non-radial. We categorized these cells as atypical astrocytes (Fig. 2E). Some of these atypical astrocytes co-expressed Nestin, while others co-expressed S100β.

Quantitative analysis using a two-way ANOVA showed a main effect of sex for the number of EYFP+ cells in animals treated from P0-P7 and from P14-P21 (Table S4), so male and female animals were separated for Nestin lineage analysis. Despite the main effect of sex, we found similar effects of treatment across sex in mice treated with VGCV for both treatment windows, indicating that sex does not determine the long-term response to VGCV. P0-P7 VGCV treated Tk+ males and females had fewer EYFP+ cells than Tk-controls (males: t_(6)_ = 6.836, p=0.0005; females: t_(6)_ = 4.917, p=0.0027), indicating that there were fewer cells in the adult Nestin lineage following neonatal VGCV treatment (Fig. 3C, E). Administering VGCV from P0-P7 led to an almost complete absence of DCX and/or NeuN expressing neurons (males: t(6) = 11.51, p<0.0001; females: t_(6)_ = 7.818, p=0.0002), but did not change the number of GFAP+ astrocytes (males: t_(6)_ = 1.014, p=0.3497; females: t_(6)_ = 0.2606, p=0.8031) (Fig. S2B, C) within the Nestin lineage in both Tk+ males and females compared to Tk-littermate controls. Since P0-P7 VGCV led to a dramatic reduction in the number of neurons and no change in the number of astrocytes in the Nestin lineage of Tk+ animals, the percentage of DCX and NeuN expressing neurons within the lineage was accordingly diminished (males: t_(6)_ = 18.72, p<0.0001; females: t_(6)_ = 20.71, p<0.0001) and the percentage of GFAP expressing astrocytes was proportionately increased (males: t_(6)_ = 21.04, p<0.0001; females: t_(6)_ = 16.35, p<0.0001) in both Tk+ males and females compared to Tk-mice (Fig. S2D, E). We have previously validated this quadruple labelling approach to account for over 90% of cells in the Nestin lineage (Dranovsky et al., 2011). The few unidentified cells (DCX-NeuN-GFAP-) may be intermediate progenitors that do not express any of these markers or may be unlabeled due to technical limitations of slice immunohistochemistry. Subpopulation assessment of immature and mature neurons demonstrated that P0-P7 VGCV resulted in very few adult-born immature (males: t_(6)_ = 12.22, p<0.0001; females: t_(6)_ = 9.001, p=0.0001) and mature (males: t_(6)_ = 8.592, p=0.0001; females: t_(6)_ = 4.833, p=0.0029) neurons in the Nestin lineage of Tk+ males and females compared to Tk-controls (Fig. 3D, F). Importantly, DCX+ cells were nearly absent throughout the dorsal-ventral DG axis in both male and female Tk+ animals. These results demonstrate that ablation of proliferating neural stem cells during the first week of life nearly abolished adult DG neurogenesis, but not adult gliogenesis.

We next examined the effects of early life VGCV on astrocytes produced by adult stem cells. We quantified the number of EYFP+GFAP+ RGLs, stellate astrocytes, and atypical astrocytes as defined above to determine the effect of VGCV treatment on these cell types in the Nestin lineage. P0-P7 VGCV led to a significant reduction in the number of RGLs (males: t_(6)_ = 5.208, p=0.002; females: t_(6)_ = 8.272, p=0.0002) and an increase in the number of atypical astrocytes (males: t_(6)_ = 4.288, p=0.0052; females: t_(6)_ = 2.644, p=0.0383) within the adult Nestin lineage in Tk+ males and females compared to Tk-animals, but the number of stellate astrocytes in the lineage was not affected in either sex (males: t_(6)_ = 0.2840, p=0.786; females: t_(6)_ = 0.5339, p=0.6126) (Fig. 3G, H). These data indicate that elimination of dividing neural stem cells during the first postnatal week resulted in alterations in RGL and atypical astrocyte production, but no effect on the stellate astrocyte component of the adult Nestin lineage.

Although a diminished pool of adult RGLs did remain after P0-P7 stem cell ablation, the treated mice almost entirely lacked adult neurogenesis. Moreover, morphologically atypical astrocytes constituted a considerable population in the Tk+ mice raising the possibility that a cell type with unclear stem cell potential was increasingly represented within the Nestin lineage following brief stem cell ablation. These observations raise an intriguing possibility that early life stem cell ablation compromises adult Nestin lineage astrocyte attributes and function. We therefore decided to examine the expression of stem and non-stem astrocyte markers in EYFP+GFAP+ astrocytes in our animals. We assessed for expression of Nestin, a marker of stem cells, and S100β, a marker of non-stem astrocytes, in the GFAP+ population of the Nestin lineage (Fig. 2C-E). Nestin+S100β-RGLs are the neural stem cells of the DG (Encinas et al., 2013; Ma et al., 2009). Nestin-S100β-RGLs may also function as stem cells, but the stem potential of S100β+ RGLs is unknown (Encinas et al., 2013). One study has reported that a small population of RGLs express S100β and have diminished stem cell capacity (Gebara et al., 2016). P0-P7 VGCV treatment led to a reduction in the number of Nestin-S100β-(males: t_(4)_ = 4.260, p=0.013; females: t_(4)_ = 3.882, p=0.0178) and Nestin+S100β-(males: t_(4)_ = 4.783, p=0.0088; females: t_(4)_ = 10.29, p=0.0005) RGLs in the Nestin lineage of both Tk+ males and females compared to Tk-animals of each sex (Fig. 3I, J). As expected, the number of S100β+ (including both Nestin-S100β+ and Nestin+S100β+) RGLs in the lineage was small in both Tk-males and females. No difference was detected in the number of these cells between Tk- and Tk+ males (t_(4)_ = 0.0, p>0.9999) (Fig. 3I). There was a decrease in the number of S100β+ RGLs in Tk+ compared to Tk- females (t_(4)_ = 3.182, p=0.0335), but the effect size of the reduction was very small (Fig. 3J). There was no difference detected in the percentage of Nestin-S100β-(males: t_(4)_ = 0.6235, p=0.5668; females: t_(4)_ = 0.3518, p=0.7427) and Nestin+S100β-(males: t_(4)_ = 2.452, p=0.0703; females: t_(4)_ = 0.6417, p=0.556) RGLs within the Nestin lineage between Tk- and Tk+ animals in both males and females (Fig. S2F, H), indicating that adult RGLs maintain their subpopulations following P0-P7 VGCV. Similarly, the atypical astrocyte population in the lineage contained both Nestin+ and S100β+ cells, and P0-P7 VGCV treatment did not affect the percentage of cells in this population that expressed each marker in males or females (Nestin-S100β-: males: t(4) = 0.08396, p=0.9371; females: t_(4)_ = 0.3975, p=0.7113. Nestin+S100β-: males: t_(4)_ = 0.2648, p=0.8043; females: t_(4)_ = 0.1516, p=0.8869. S100β+: males: t_(4)_ = 0.3787, p=0.7241; females: t_(4)_ = 0.06041, p=0.9547) (Fig. S2G, I). Additionally, not all stellate astrocytes within the Nestin lineage expressed S100β, but VGCV treatment did not alter the percentage of stellate astrocytes that did or did not express this marker in males or females (Nestin-S100β-: males: t_(4)_ = 0.7047, p=0.5199; females: t_(4)_ = 0.8100, p=0.4634. Nestin-S100β+: males: t_(4)_ = 0.7047, p=0.5199; females: t_(4)_ = 0.8100, p=0.4634) (Fig. S2J, K). These results demonstrate that a one-week ablation of proliferating stem cells during the first week of life leads to a lasting depletion of the adult stem cell pool in both males and females with a corresponding inhibition of adult neurogenesis and a neural stem cell fate bias towards astrocytic progeny.

Previously, we reported that elimination of dividing stem cells during early adolescence (P28-P35) did not result in lasting effects on the size of the Nestin lineage (Kirshenbaum et al., 2014). Given our findings with the P0-P7 treatment above, we sought to establish whether the sensitive period for affecting adult neurogenesis is closed by the third postnatal week, when neurogenesis begins to resemble its adult anatomical restriction (Nicola et al., 2015; Sugiyama et al., 2013). We assessed the adult Nestin lineage in mice treated with VGCV from P14-P21 (Fig. 4A, B) and found that the treatment led to fewer EYFP+ cells in both Tk+ males and females compared to Tk-animals of each sex (males: t_(10)_ = 2.365, p=0.0396; females: t_(9)_ = 4.575, p=0.0013) (Fig. 4C, E). P14-P21 VGCV led to fewer DCX and NeuN expressing neurons (males: t_(10)_ = 4.055, p=0.0023; females: t_(9)_ = 9.883, p<0.0001), but no change in the number of GFAP expressing astrocytes (males: t_(10)_ = 0.2030, p=0.8432; females: t_(9)_ = 0.4325, p=0.6756) in the Nestin lineage of Tk+ compared to Tk-mice for both males and females (Fig. S3B, C). Consequently, the percentage of neurons within the Nestin lineage was decreased (males: t_(10)_ = 7.533, p<0.0001; females: t_(9)_ = 7.659, p<0.0001), while the percentage of astrocytes within the lineage was increased (males: t_(10)_ = 6.591, p<0.0001; females: t_(9)_ = 7.743, p<0.0001) in both Tk+ males and females compared to Tk-controls (Fig. S3D, E). P14-P21 VgCV led to fewer adult-born immature (males: t_(10)_ = 4.533, p=0.0011; females: t_(9)_ = 10.72, p<0.0001) and mature (males: t_(10)_ = 2.226, p=0.0501; females: t_(9)_ = 5.412, p=0.0004) neuron subpopulations in the Nestin lineage of Tk+ compared to Tk-mice (Fig. 4D, F). The number of DCX+ cells was reduced across the dorsal-ventral axis of the DG in both Tk+ males and females. In contrast to the P0-P7 treated animals, P14-P21 treated Tk+ animals did not have a decrease in the number of RGLs in the lineage for either sex compared to Tk-animals (males: t_(10)_ = 0.7595, p=0.4651; females: t_(9)_ = 1.190, p=0.2643) (Fig. 4G, H). There was also no difference detected in the number of EYFP+ stellate (males: t_(10)_ = 0.03963, p=0.9692; females: t_(9)_ = 0.03157, p=0.9755) or atypical (males: t_(10)_ = 1.646, p=0.1308; females: t_(9)_ = 1.214, p=0.2555) astrocytes between Tk- and Tk+ animals in both sexes (Fig. 4G, H). These results indicate that ablation of proliferating stem cells during the third postnatal week leads to reduced adult neurogenesis and a bias towards astrocytic progeny.

Analysis of stem and non-stem marker expression in GFAP+ astrocytes in the Nestin lineage demonstrated that there were no differences in the number of Nestin-S100β-(males: t_(6)_ = 0.1491, p=0.8864; females: t_(6)_ = 1.254, p=0.2564), Nestin+S100β-(males: t_(6)_ = 1.310, p=0.2382; females: t_(6)_ = 0.6512, p=0.539) or S100β+ RGLs (males: t_(6)_ = 0.7146, p=0.5017; females: t_(6)_ = 0.4763, p=0.6507) in Tk+ compared to Tk-mice (Fig. 4I, J). Moreover, no differences were detected in the percentage of RGLs expressing these markers (Nestin-S100β-: males: t_(6)_ = 0.4798, p=0.6483; females: t_(6)_ = 1.641, p=0.1519. Nestin+S100β-: males: t_(6)_ = 0.3597, p=0.7314; females: t_(6)_ = 1.662, p=0.1476. S100β+: males: t_(6)_ = 0.6074, p=0.5658; females: t_(6)_ = 0.5694, p=0.5898) (Fig. S3F, H), indicating that P14-P21 VGCV treatment did not alter the stem and non-stem marker expression of these cells or decrease the pool of adult neural stem cells. While atypical astrocytes reflected a minor component of the GFAP+ Nestin lineage in this experiment, both Nestin+ and S100β+ cells were represented. P14-P21 VGCV treatment did not alter the percentage of cells expressing these markers in male Tk+ compared to Tk-animals (Nestin-S100β-: t_(6)_ = 0.9019, p=0.4019; Nestin+S100β-: t_(6)_ = 0.8662, p=0.4197; S100β+: t_(6)_ = 1.015, p=0.3492) (Fig. S3G). Tk+ P14-P21 treated females had an increase in the percentage of Nestin+S100β-cells within the atypical astrocytes (t_(6)_ = 2.756, p=0.0330) compared to Tk-females, but no difference was detected in the percentage of Nestin-S100β-(t_(6)_ = 1.500, p=0.1842) or S100β+ cells in this population (t_(6)_ = 0.4973, p=0.6367) (Fig. S3I). Unlike following P0-P7 VGCV, however, the total number of atypical astrocytes was small in all P14-P21 treated groups (Fig. 4G, H). Lastly, VGCV treatment did not alter the percentage of stellate astrocytes in the lineage that did or did not express S100β+ in males or females (Nestin-S100β-: males: t_(6)_ = 1.005, p=0.3538; females: t_(6)_ = 0.7869, p=0.4613. Nestin-S100β+: males: t_(6)_ = 1.005, p=0.3538; females: t_(6)_ = 0.7869, p=0.4613) (Fig. S3J, K).Together, these results demonstrate that P14-P21 VGCV treatment in males and females reduced adult neurogenesis and created a neural stem cell fate bias towards astrocytic progeny, and that these lineage changes do not simply reflect a depleted neural stem cell pool as the number of adult RGLs was maintained.

### Neural stem cell proliferation during early life

Our results show that VGCV treatment during the first, but not the third, week of life leads to fewer adult stem cells within the Nestin lineage. Since stem cells present in early life are thought to be the source of the adult stem cell pool (Encinas et al., 2013; Li et al., 2013; Nicola et al., 2015; Sugiyama et al., 2013), the P0-P7 time period may be susceptible to a life-long decrease in RGLs due to the greater number of dividing GFAP+ stem cells available for VGCV targeting during this time. Indeed, postnatal day 6, compared to P3 or P9, has been suggested to be the peak of proliferation for stem cells that remain into adulthood (Ortega-Martinez and Trejo, 2015). Therefore, we hypothesized that there are more dividing stem cells during the P0-P7 period compared to the P14-P21 period, and that this difference may underlie the ability of VGCV treatment to reduce the adult stem cell pool after treatment during the earlier time window.

To test this hypothesis directly, we used a transgenic mouse line with expression of a rapidly degrading fluorescent Kusabira Orange (KOr) protein under the control of a Nestin promoter (Ishizuka et al., 2011; Kanki et al., 2010) to visualize DG stem cells in animals sacrificed at P7 or P21 (Fig. 5A). Qualitative assessment suggested that there are many more KOr+ cells with radial processes in the developing DG at P7 than at P21, supporting the notion that more RGL stem cells are present at P7 versus P21 (Fig. 5B). We wanted to quantify the percentage of stem cells dividing at each time point that would be targeted by VGCV in our GFAP-Tk mouse line. We counted the number of KOr+ cells that expressed GFAP in their radial processes, then assessed for co-expression of MCM2, a cell division marker. Two-way ANOVA analysis with sex and age as fixed factors identified no main effect of sex (F_(1,8)_ = 0.8343, p=0.3877) or interaction of sex x age (F_(1,8)_ = 2.031, p=0.1919), but there was a main effect of age (F_(1,8)_ = 608.1, p<0.0001), with a greater percentage of KOr+GFAP+ cells expressing MCM2 at P7 versus at P21 (Fig. 5C). These results demonstrate that there are more dividing GFAP+ stem cells at P7 compared to P21. These data suggest that the long-term effect on stem cells is more severe with treatment from P0-P7 compared to P14-P21 likely due to greater proliferation, and thus ablation, of stem cells earlier in life.

**Figure 5.**
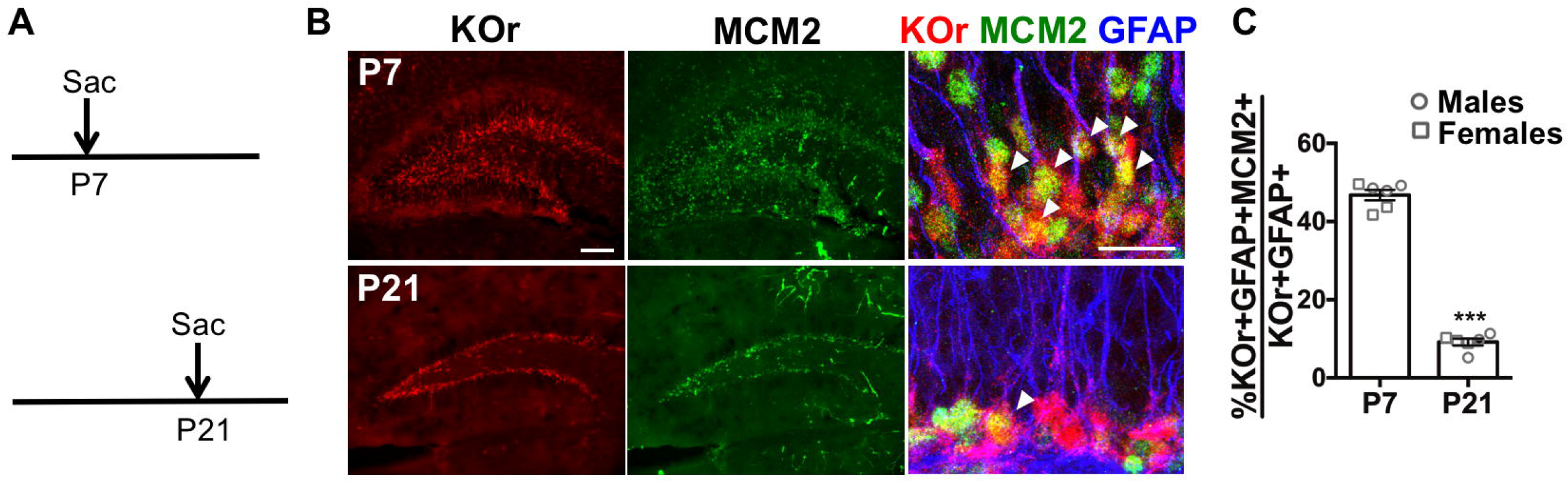
More DG stem cells are dividing during the first versus third postnatal week. (A) Experimental timeline of sacrifice of Nestin-Kusabira Orange (KOr) animals at P7 or P21. (B) Representative images of KOr+, MCM2+, and GFAP+ cells in P7 and P21 mice. Arrowheads point to cells expressing all three markers. (C) A larger percentage of KOr+GFAP+ RGLs express the cell division marker MCM2 at P7 compared to at P21 in males and females. Scale bars represent 100 μM for low magnification and 20 μM for high magnification images. Data are expressed as mean ± SEM. ***p<0.001

## Discussion

### The first three postnatal weeks constitute sensitive periods for determining adult neural stem cell number and lineage potential

In this study, we asked whether ablation of proliferating neural stem cells during early life is sufficient to lead to long-term changes in adult stem lineage potential. Our results suggest that the first three weeks of life constitute a series of sensitive periods during which targeting proliferating DG stem cells can have lasting effects on adult stem cell lineages. Importantly, the first postnatal week constitutes a sensitive window for establishing an adult stem cell pool. During this time, ablating dividing stem cells depletes the adult stem cell pool and expands a population of astrocytes that have unclear stem potential. By the third postnatal week, the stem cell pool appears formed, as ablation of proliferating neural stem cells during this period does not affect the number of stem cells present in the adult lineage. These results indicate that the sensitive window for determining the size of the adult stem cell pool is closed by the third postnatal week. Both interventions, however, reduce the number of cells in the adult stem cell lineage, decrease adult neurogenesis, and produce a lineage bias toward astrocytic progeny. We previously reported that ablation of proliferating neural stem cells during the fifth postnatal week does not produce lasting effects on the size of the Nestin stem cell lineage (Kirshenbaum et al., 2014). In this context, our data demonstrating that ablation of dividing stem cells during the first or third postnatal week leads to smaller adult Nestin lineages suggests that the sensitive window for affecting the size of the adult stem cell lineage is closed by the fifth postnatal week. The absence of effect on the Nestin lineage size after fifth postnatal week stem cell ablation suggests that adult neurogenesis may not be altered when dividing stem cells are targeted during this time period, but future studies are needed to further explore this possibility. Important to note, however, is that the severity of stem cell ablation, determined by the VGCV dose and duration of treatment, likely interacts with the timing of treatment to determine the long-term effects, so a more severe adult ablation could potentially produce persisting effects. Our findings demonstrate that transient elimination of dividing stem cells during early life is sufficient to lead to persistent changes in adult stem cell lineage specification and underscore the first and third postnatal weeks as distinct sensitive periods for the effects of ablating dividing stem cells on the adult stem cell lineage. These data parallel the results of studies investigating the long-term effects of drug administration or chronic stress at different times during life: Interventions during the first three weeks of life lead to lasting deficits on adult DG cell proliferation and neurogenesis (Aisa et al., 2009; Hulshof et al., 2011; Kikusui and Mori, 2009; Leslie et al., 2011; Mirescu et al., 2004; Naninck et al., 2015; Suri et al., 2013; Yu et al., 2010; Yu et al., 2017b; Zhu et al., 2010), while the effects of similar interventions during adolescence or adulthood recover after the intervention is terminated (Elizalde et al., 2010; Heine et al., 2004; Lagace et al., 2010; Zhu et al., 2010).

Sensitive developmental windows for determining stem cell function or response to environmental insults have been previously demonstrated for neural stem cells in multiple brain areas. DG stem and progenitor cells have been shown to be sensitive to the anti-proliferative and anti-neurogenic effects of the environmental toxicant methylmercury at P7 and P14, but not P21 (Obiorah et al., 2015). The temporal restriction of the negative effects of methylmercury to earlier developmental time periods is analogous to the more robust and long-term effects of VGCV treatment earlier in life. Additionally, in normal cortical development, stem cell function changes as the brain assembles. During early cortical development, radial cells predominantly divide symmetrically to expand the stem cell pool, whereas later in development, these cortical stem cells more often undergo differentiating divisions to give rise to neurons (Gotz and Huttner, 2005). These examples underscore how stem cell lineage specification changes as the brain matures. Hence, interfering with the DG neural stem pool during distinct periods during early development may have distinct outcomes on stem cell lineage trajectories for the life of the animal. In the case of DG neural stem cells, which are exquisitely sensitive to the effects of chronic stress and other environmental insults, these sensitive periods may determine the longterm effects of these insults on neurogenesis.

### Early life ablation of proliferating stem cells alters adult stem cell lineage specification

Our first analysis of the DG Nestin lineage showed that ablating dividing stem cells from P0-P7 results in a 50% reduction in the number of RGL stem cells, but almost no neurogenesis within the adult Nestin lineage. Fewer adult stem cells likely reflect direct depletion of young RGLs due to the intervention since more stem cells dividing during the first postnatal week make RGLs an especially vulnerable population during this time. However, the near total reduction in neurogenesis far outweighs the 50% decrease in stem cells, suggesting that the remaining stem cells have diminished capacity for neurogenesis within their stem cell niche. Observations that targeting dividing stem cells from P14-P21 does not affect the number of RGLs in the adult lineage, but still leads to a marked reduction in neuronal progeny further support the notion that ablation of stem cells during early life appears to alter the neurogenic capacity of adult stem cells. Conversely, early life dividing stem cell ablation does not decrease and may even increase gliogenesis, as equal numbers of stellate astrocytes are present in the adult Nestin lineage despite fewer adult stem cells after P0-P7 treatment. These findings suggest that either stellate astrocyte production or survival is increased to maintain the population.

We explored a potential stem cell-autonomous explanation for diminished neurogenesis in the adult animal by assessing for stem and non-stem markers in RGLs. A decrease in the percentage of RGLs expressing stem marker Nestin or an increase in the number of RGLs expressing the non-stem marker S100β would have suggested diminished stem cell potential of RGLs with a more terminal astrocyte-like phenotype (Gebara et al., 2016), but our results do not support this possibility. Our data demonstrate that RGLs in Tk+ animals are still functional in their capacity to self-renew and to make stellate astrocytes, but their neurogenic capacity is decreased. These RGLs may be intrinsically unable to produce the intermediate progenitors that give rise to neuroblasts or the DG neurogenic niche, in which RGLs and other GFAP+ non-stem astrocytes are thought to play a role (Alvarez-Buylla and Lim, 2004), may be unable to promote the differentiation and/or survival of neuroblasts and newborn neurons. Thus, the long-term effects of VGCV treatment on the Nestin lineage may be the result of cell autonomous changes in the adult RGLs or secondary consequences of a stem cell niche that has been compromised in response to depletion of stem cells and/or possibly non-stem stellate astrocytes. Our results suggest the immediate recovery of cell proliferation following removal of VGCV in early life followed by decreased neurogenesis in adulthood. This would be more likely to occur via an evolving change in stem cell niche function, though the possibility of a slowly emerging autonomous decline of stem cell potential cannot be ruled out.

Both stem cell-autonomous and niche effects have been shown to contribute to changes in DG cell proliferation and neurogenesis in response to the environment or interventions. For instance, fingolimod, an immune response-mediating drug, increases the proliferation and survival of DG stem cells and their neuronal progeny in vivo and in vitro, indicating that the effect of this drug on stem cell activity is cell-autonomous (Efstathopoulos et al., 2015). In contrast, both voluntary wheel running and short-term calorie restriction alter molecular niche factors to promote DG neurogenesis without impacting the number of stem cells, demonstrating that neurogenesis can be modulated through environmentally-induced niche effects (Dranovsky et al., 2011; Gremmelspacher et al., 2017; Hornsby et al., 2016). Interestingly, the long-term effects of early life ablation of dividing stem cells are similar to the effects of hippocampal irradiation, suggesting that the mechanisms underlying these responses may also be analogous. As with early life dividing stem cell ablation, irradiation either during early life or adulthood results in dramatically diminished DG neurogenesis, but maintenance of the stem/progenitor cell pool and preserved gliogenesis (Dranovsky et al., 2011; Mizumatsu et al., 2003; Monje et al., 2002; Naylor et al., 2008; Rola et al., 2004). Decreased neurogenesis seen after irradiation is hypothesized to be due mainly to an unsupportive niche (Dranovsky et al., 2011; Monje et al., 2002). The potential long-term term effects of early life stem cell ablation on DG stem cell-autonomous activity and the stem cell niche, as well as the mechanisms by which these effects occur, require further investigation.

### Sex as a variable largely does not contribute to determining response to VGCV

The effects of VGCV treatment on the adult Nestin lineage are almost indistinguishable between male and female animals for each treatment window, indicating that sex does not contribute to determining the long-term effects of VGCV treatment. However, we did detect a main effect of sex in the number of cells in the Nestin lineage, with more cells in the lineage of female animals. Previous studies using similar methods of lineage tracing found that there are more cells in the Nestin lineage of female compared to male animals 12 days after induction of labeling, but the difference was not statistically significant (Lagace et al., 2007). In this context, a significant difference in the size of the lineages between sexes 7 weeks after initiation of labeling suggests that the disparity in cell number is accentuated as the lineage accumulates. The larger lineages of female animals could be because 1) there are more Nestin+ stem and progenitor cells available to be labelled initially in females, 2) Nestin+ stem and progenitor cells are more active and produce larger lineages in females, and/or 3) TMX metabolism differs between males and females so that more stem and progenitor cells are initially labelled in females. Regardless of these possibilities, the proportions of neurons and astrocytes in the Nestin lineage of males and females in each treatment group are similar, suggesting similar lineage potentials of DG RGLs between the sexes.

We also found that P0-P7 VGCV females and males differ in the recovery of cell proliferation one week after treatment completion. However, the adult effects of P0-P7 VGCV treatment are almost identical in males and females, suggesting that the difference in short-term recovery of cell proliferation does not translate to differences in long-term effects and that the 1-2 week suppression of stem cell proliferation during early life is sufficient to produce the long-term effects in both sexes. The continued suppression of cell proliferation in female mice observed one week after P0-P7 VGCV suggests that the stem cells or neurogenic niche in females may be more sensitive in the short-term to stem cell ablation at this early age.

### Stem cell ablation and reduction in cell proliferation may contribute to persistent effects of early life insults

Unlike with exposure during adulthood, early life exposures to social and chemical stress in humans have a propensity to result in lasting adverse effects on brain function, potentially inducing a variety of neurological and psychiatric conditions that include deficits in hippocampal function (Cooper et al., 2015; Dorn et al., 2014; Rees and Inder, 2005). These observations parallel persisting hippocampal sequelae of neonatal and juvenile exposure of mice to drugs, infections and psychological stress (McEwen, 2003; Ruddy and Morshead, 2018; Yu et al., 2017a). Curiously, these early life manipulations in mice lead to lasting deficits in DG stem cell number and neurogenesis, but the mechanisms by which these persistent changes develop are unknown. In this study, we utilized early life ablation of dividing stem cells to mimic the stem cell depletion, diminished cell proliferation and neurogenesis, and increased apoptosis that occurs immediately or shortly after biological early life insults. We demonstrate that this early postnatal stem cell ablation is sufficient to deplete the adult stem cell pool and decrease adult neurogenesis. Thus, these short-term cellular effects of early postnatal insults have the capacity to produce the long-term DG stem cell lineage deficits. The possibility that these life-long changes in DG neurogenesis and lineage homeostasis contribute to persisting hippocampal dysfunction associated with early life stress remains to be explored.

## Acknowledgements

The authors thank Laura Checkley and Jennifer Payne for technical assistance, and Míriam Navarro-Sobrino, Lillian Lawrence and Alvaro Garcia-Garcia for critical reading of the manuscript. We also thank Hiroaki Kanki and Hideyuki Okano for providing Nestin-Kusabira Orange mice. This work was supported by MH106809, MH091844, Columbia University Irving Scholar Award (AD) and MH105675 (EdL). MY was supported by F30MH111209 and T32GM007367, GSK was supported by a Canadian Institutes of Health Research Postdoctoral Fellowship, and PA was supported by Veni Grant 016.161.045.

---

**Figure S1. Schematic hierarchical categorization of cells within the Nestin lineage.**

**Figure S2. P0-P7 VGCV alters the neurogenic and astrocytic adult Nestin lineage potential, without affecting astrocyte stem and non-stem marker expression.** (A) Experimental timeline of P0-P7 VGCV treatment and TMX administration in GFAP-Tk/Nestin-CreER^T2^ mice. (B) P0-P7 VGCV reduced the number of DCX+ and/or NeuN+ neurons, but did not change the number of GFAP+ astrocytes, within the Nestin lineage of Tk+ males and (C) females compared to Tk-animals. (D) P0-P7 VGCV decreased the percentage of DCX and/or NeuN expressing neurons and increased the percentage of GFAP expressing astrocytes, but did not affect the percentage of unidentified DCX-NeuN-GFAP-cells within the Nestin lineage of Tk+ males and (E) females versus Tk-animals. (F) No difference was detected in the percentage of Nestin-S100β- or Nestin+S100β-RGLs, but there was an increase in the percentage of S100β+ RGLs in Tk+ compared to Tk-males. (G) No difference was detected in the percentage of Nestin-S100β-, Nestin+S100β-, or S100β+ atypical astrocytes in P0-P7 VGCV Tk-versus Tk+ males. (H) No difference was detected in the percentage of Nestin-S100β-, Nestin+S100β-, S100β+ RGLs or (I) atypical astrocytes between P0-P7 VGCV Tk-and Tk+ females. (J) No difference was detected in the percentage of Nestin-S100β- or Nestin-S100β+ stellate astrocytes in P0-P7 VGCV Tk-versus Tk+ males or (K) females. Data are expressed as mean ± SEM. *p≤0.05, ***p<0.001

**Figure S3. P14-P21 VGCV alters the neurogenic and astrocytic adult Nestin lineage potential, without affecting astrocyte stem and non-stem marker expression.** (A) Experimental timeline of P14-P21 VGCV treatment and TMX administration in GFAP-Tk/Nestin-CreER^T2^ mice. (B) P14-P21 VGCV reduced the number of DCX+ and/or NeuN+ neurons, but did not change the number of GFAP+ astrocytes, within the Nestin lineage of Tk+ males and (C) females compared to Tk-animals. (D) P14-P21 VGCV decreased the percentage of DCX and/or NeuN expressing neurons and increased the percentage of GFAP expressing astrocytes and DCX-NeuN-GFAP-unidentified cells within the Nestin lineage of Tk+ males and (E) females versus Tk-animals. (F) No difference was detected in the percentage of Nestin-S100β-, Nestin+S100β-, or S100β+ RGLs or (G) atypical astrocytes in P14-P21 VGCV Tk-versus Tk+ males. (H) No difference was detected in the percentage of Nestin-S100β-, Nestin+S100β-, S100β+ RGLs between P14-P21 VGCV Tk- and Tk+ females. (I) P14-P21 VGCV increased the percentage of Nestin+S100β-, but did not affect the percentage of Nestin-S100β- or S100β+, atypical astrocytes in Tk+ versus Tk-females. (J) No difference was detected in the percentage of Nestin-S100β- or Nestin-S100β+ stellate astrocytes in P14-P21 VGCV Tk-versus Tk+ males or (K) females. Data are expressed as mean ± SEM. *p≤0.05, **p<0.01, ***p<0.001

## References

Aisa B, Elizalde N, Tordera R, Lasheras B, Del Rio J, Ramirez MJ. 2009. Effects of neonatal stress on markers of synaptic plasticity in the hippocampus: implications for spatial memory. Hippocampus 19(12):1222–31.

Altman J, Bayer SA. 1990. Migration and distribution of two populations of hippocampal granule cell precursors during the perinatal and postnatal periods. J Comp Neurol 301(3):365–81.

Alvarez-Buylla A, Lim DA. 2004. For the long run: maintaining germinal niches in the adult brain. Neuron 41(5):683–6.

Angevine JB, Jr. 1965. Time of neuron origin in the hippocampal region. An autoradiographic study in the mouse. Exp Neurol Suppl:Suppl 2:1–70.

Baek SB, Bahn G, Moon SJ, Lee J, Kim KH, Ko IG, Kim SE, Sung YH, Kim BK, Kim TS and others. 2011. The phosphodiesterase type-5 inhibitor, tadalafil, improves depressive symptoms, ameliorates memory impairment, as well as suppresses apoptosis and enhances cell proliferation in the hippocampus of maternal-separated rat pups. Neurosci Lett 488(1):26–30.

Baek SS, Jun TW, Kim KJ, Shin MS, Kang SY, Kim CJ. 2012. Effects of postnatal treadmill exercise on apoptotic neuronal cell death and cell proliferation of maternal-separated rat pups. Brain Dev 34(1):45–56.

Bayer SA. 1980. Development of the hippocampal region in the rat. I. Neurogenesis examined with 3H-thymidine autoradiography. J Comp Neurol 190(1):87–114.

Bayer SA, Altman J. 1974. Hippocampal development in the rat: cytogenesis and morphogenesis examined with autoradiography and low-level X-irradiation. J Comp Neurol 158(1):55–79.

Bonaguidi MA, Wheeler MA, Shapiro JS, Stadel RP, Sun GJ, Ming GL, Song H. 2011. In vivo clonal analysis reveals self-renewing and multipotent adult neural stem cell characteristics. Cell 145(7):1142–55.

Brown JP, Couillard-Despres S, Cooper-Kuhn CM, Winkler J, Aigner L, Kuhn HG. 2003. Transient expression of doublecortin during adult neurogenesis. J Comp Neurol 467(1):1–10.

Bush TG, Savidge TC, Freeman TC, Cox HJ, Campbell EA, Mucke L, Johnson MH, Sofroniew MV. 1998. Fulminant jejuno-ileitis following ablation of enteric glia in adult transgenic mice. Cell 93(2):189–201.

Cameron HA, Glover LR. 2015. Adult neurogenesis: beyond learning and memory. Annu Rev Psychol 66:53–81.

Cooper jM, Gadian DG, Jentschke S, Goldman A, Munoz M, Pitts G, Banks T, Chong WK, Hoskote A, Deanfield J and others. 2015. Neonatal hypoxia, hippocampal atrophy, and memory impairment: evidence of a causal sequence. Cereb Cortex 25(6):1469–76.

Dorn M, Lidzba K, Bevot A, Goelz R, Hauser TK, Wilke M. 2014. Long-term neurobiological consequences of early postnatal hCMV-infection in former preterms: a functional MRI study. Hum Brain Mapp 35(6):2594–606.

Dranovsky A, Picchini AM, Moadel T, Sisti AC, Yamada A, Kimura S, Leonardo ED, Hen R. 2011. Experience dictates stem cell fate in the adult hippocampus. Neuron 70(5):908–23.

Efstathopoulos P, Kourgiantaki A, Karali K, Sidiropoulou K, Margioris AN, Gravanis A, Charalampopoulos I. 2015. Fingolimod induces neurogenesis in adult mouse hippocampus and improves contextual fear memory. Transl Psychiatry 5:e685.

Elizalde N, Garcia-Garcia aL, Totterdell S, Gendive N, Venzala E, Ramirez MJ, Del Rio J, Tordera RM. 2010. Sustained stress-induced changes in mice as a model for chronic depression. Psychopharmacology (Berl) 210(3):393–406.

Encinas JM, Sierra A, Valcarcel-Martin R, Martin-Suarez S. 2013. A developmental perspective on adult hippocampal neurogenesis. Int J Dev Neurosci 31(7):640–5.

Fantetti KN, Gray EL, Ganesan P, Kulkarni A, O’Donnell LA. 2016. Interferon gamma protects neonatal neural stem/progenitor cells during measles virus infection of the brain. J Neuroinflammation 13(1):107.

Garcia AD, Doan NB, Imura T, Bush TG, Sofroniew MV. 2004. GFAP-expressing progenitors are the principal source of constitutive neurogenesis in adult mouse forebrain. Nat Neurosci 7(11):1233–41.

Gebara E, Bonaguidi MA, Beckervordersandforth R, Sultan S, Udry F, Gijs PJ, Lie DC, Ming GL, Song H, Toni N. 2016. Heterogeneity of Radial Glia-Like Cells in the Adult Hippocampus. Stem Cells 34(4):997–1010.

Gotz M, Huttner Wb. 2005. The cell biology of neurogenesis. Nat Rev Mol Cell Biol 6(10):777–88.

Gremmelspacher T, Gerlach J, Hubbe A, Haas CA, Haussler U. 2017. Neurogenic Processes Are Induced by Very Short Periods of Voluntary Wheel-Running in Male Mice. Front Neurosci 11:385.

Heine VM, Maslam S, Zareno J, Joels M, Lucassen PJ. 2004. Suppressed proliferation and apoptotic changes in the rat dentate gyrus after acute and chronic stress are reversible. Eur J Neurosci 19(1):131–44.

Hornsby AK, Redhead YT, Rees DJ, Ratcliff MS, Reichenbach A, Wells T, Francis L, Amstalden K, Andrews ZB, Davies JS. 2016. Short-term calorie restriction enhances adult hippocampal neurogenesis and remote fear memory in a Ghsr-dependent manner. Psychoneuroendocrinology 63:198–207.

Hulshof hJ, Novati A, Sgoifo A, Luiten PG, den Boer JA, Meerlo P. 2011. Maternal separation decreases adult hippocampal cell proliferation and impairs cognitive performance but has little effect on stress sensitivity and anxiety in adult Wistar rats. Behav Brain Res 216(2):552–60.

Ishizuka K, Kamiya A, Oh EC, Kanki H, Seshadri S, Robinson JF, Murdoch H, Dunlop AJ, Kubo K, Furukori K and others. 2011. DISC1-dependent switch from progenitor proliferation to migration in the developing cortex. Nature 473(7345):92–6.

Kalm M, Fukuda A, Fukuda H, Ohrfelt A, Lannering B, Bjork-Eriksson T, Blennow K, Marky I, Blomgren K. 2009. Transient inflammation in neurogenic regions after irradiation of the developing brain. Radiat Res 171(1):66–76.

Kanki H, Shimabukuro MK, Miyawaki A, Okano H. 2010. “Color Timer” mice: visualization of neuronal differentiation with fluorescent proteins. Mol Brain 3:5.

Kikusui T, Mori Y. 2009. Behavioural and neurochemical consequences of early weaning in rodents. J Neuroendocrinol 21(4):427–31.

Kirshenbaum GS, Lieberman SR, Briner TJ, Leonardo ED, Dranovsky A. 2014. Adolescent but not adult-born neurons are critical for susceptibility to chronic social defeat. Front Behav Neurosci 8:289.

Kuhn GH, Blomgren K. 2011. Developmental dysregulation of adult neurogenesis. Eur J Neurosci 33(6):1115–22.

Lagace DC, Donovan MH, DeCarolis NA, Farnbauch LA, Malhotra S, Berton O, Nestler EJ, Krishnan V, Eisch AJ. 2010. Adult hippocampal neurogenesis is functionally important for stress-induced social avoidance. Proc Natl Acad Sci U S A 107(9):4436–41.

Lagace DC, Whitman MC, Noonan MA, Ables JL, DeCarolis NA, Arguello aA, Donovan MH, Fischer SJ, Farnbauch LA, Beech RD and others. 2007. Dynamic contribution of nestin-expressing stem cells to adult neurogenesis. J Neurosci 27(46):12623–9.

Lajud N, Roque A, Cajero M, Gutierrez-Ospina G, Torner L. 2012. Periodic maternal separation decreases hippocampal neurogenesis without affecting basal corticosterone during the stress hyporesponsive period, but alters HPA axis and coping behavior in adulthood. Psychoneuroendocrinology 37(3):410–20.

Leslie AT, Akers KG, Krakowski AD, Stone SS, Sakaguchi M, Arruda-Carvalho M, Frankland PW. 2011. Impact of early adverse experience on complexity of adult-generated neurons. Transl Psychiatry 1:e35.

Li G, Fang L, Fernandez G, Pleasure SJ. 2013. The ventral hippocampus is the embryonic origin for adult neural stem cells in the dentate gyrus. Neuron 78(4):658–72.

Lu Y, Giri PK, Lei S, Zheng J, Li W, Wang N, Chen X, Lu H, Zuo Z, Liu Y and others. 2017. Pretreatment with minocycline restores neurogenesis in the subventricular zone and subgranular zone of the hippocampus after ketamine exposure in neonatal rats. Neuroscience 352:144–154.

Ma DK, Bonaguidi MA, Ming GL, Song H. 2009. Adult neural stem cells in the mammalian central nervous system. Cell Res 19(6):672–82.

Mathews EA, Morgenstern NA, Piatti VC, Zhao C, Jessberger S, Schinder AF, Gage FH. 2010. A distinctive layering pattern of mouse dentate granule cells is generated by developmental and adult neurogenesis. J Comp Neurol 518(22):4479–90.

McEwen BS. 2003. Early life influences on life-long patterns of behavior and health. Ment Retard Dev Disabil Res Rev 9(3):149–54.

Ming GL, Song H. 2005. Adult neurogenesis in the mammalian central nervous system. Annu Rev Neurosci 28:223–50.

Mirescu C, Peters JD, Gould E. 2004. Early life experience alters response of adult neurogenesis to stress. Nat Neurosci 7(8):841–6.

Mizumatsu S, Monje ML, Morhardt DR, Rola R, Palmer TD, Fike JR. 2003. Extreme sensitivity of adult neurogenesis to low doses of X-irradiation. Cancer Res 63(14):4021–7.

Monje ML, Mizumatsu S, Fike JR, Palmer TD. 2002. Irradiation induces neural precursor-cell dysfunction. Nat Med 8(9):955–62.

Muramatsu R, Ikegaya Y, Matsuki N, Koyama R. 2007. Neonatally born granule cells numerically dominate adult mice dentate gyrus. Neuroscience 148(3):593–8.

Naninck EF, Hoeijmakers L, Kakava-Georgiadou N, Meesters A, Lazic SE, Lucassen PJ, Korosi A. 2015. Chronic early life stress alters developmental and adult neurogenesis and impairs cognitive function in mice. Hippocampus 25(3):309–28.

Navarro-Quiroga I, Hernandez-Valdes M, Lin SL, Naegele JR. 2006. Postnatal cellular contributions of the hippocampus subventricular zone to the dentate gyrus, corpus callosum, fimbria, and cerebral cortex. J Comp Neurol 497(5):833–45.

Naylor AS, Bull C, Nilsson MK, Zhu C, Bjork-Eriksson T, Eriksson pS, Blomgren K, Kuhn HG. 2008. Voluntary running rescues adult hippocampal neurogenesis after irradiation of the young mouse brain. Proc Natl Acad Sci U S A 105(38):14632–7.

Nicola Z, Fabel K, Kempermann G. 2015. Development of the adult neurogenic niche in the hippocampus of mice. Front Neuroanat 9:53.

Obiorah M, McCandlish E, Buckley B, DiCicco-Bloom E. 2015. Hippocampal developmental vulnerability to methylmercury extends into prepubescence. Front Neurosci 9:150.

Oreland S, Nylander I, Pickering C. 2010. Prolonged maternal separation decreases granule cell number in the dentate gyrus of 3-week-old male rats. Int J Dev Neurosci 28(2):139–44.

Ortega-Martinez S, Trejo JL. 2015. The postnatal origin of adult neural stem cells and the effects of glucocorticoids on their genesis. Behav Brain Res 279:166–76.

Patrylo PR, Nowakowski RS. 1994. Morphology and distribution of astrocytes in the molecular layer of the dentate gyrus in NZB/BlNJ, Dreher, and C57BL/6J mice. Glia 10(1):1–9.

Rees S, Inder T. 2005. Fetal and neonatal origins of altered brain development. Early Hum Dev 81(9):753–61.

Rola R, Raber J, Rizk A, Otsuka S, VandenBerg SR, Morhardt DR, Fike JR. 2004. Radiation-induced impairment of hippocampal neurogenesis is associated with cognitive deficits in young mice. Exp Neurol 188(2):316–30.

Ruddy RM, Morshead CM. 2018. Home sweet home: the neural stem cell niche throughout development and after injury. Cell Tissue Res 371(1):125–141.

Seri B, Garcia-Verdugo JM, Collado-Morente L, McEwen bS, Alvarez-Buylla A. 2004. Cell types, lineage, and architecture of the germinal zone in the adult dentate gyrus. J Comp Neurol 478(4):359–78.

Srinivas S, Watanabe T, Lin CS, William CM, Tanabe Y, Jessell TM, Costantini F. 2001. Cre reporter strains produced by targeted insertion of EYFP and ECFP into the ROSA26 locus. BMC Dev Biol 1:4.

Stanfield BB, Cowan WM. 1979. The development of the hippocampus and dentate gyrus in normal and reeler mice. J Comp Neurol 185(3):423–59.

Sugiyama T, Osumi N, Katsuyama Y. 2013. The germinal matrices in the developing dentate gyrus are composed of neuronal progenitors at distinct differentiation stages. Dev Dyn 242(12):1442–53.

Suri D, Veenit V, Sarkar A, Thiagarajan D, Kumar A, Nestler EJ, Galande S, Vaidya VA. 2013. Early stress evokes age-dependent biphasic changes in hippocampal neurogenesis, BDNF expression, and cognition. Biol Psychiatry 73(7):658–66.

Xu L, Yang Y, Gao L, Zhao J, Cai Y, Huang J, Jing S, Bao X, Wang Y, Gao J and others. 2015. Protective effects of resveratrol on the inhibition of hippocampal neurogenesis induced by ethanol during early postnatal life. Biochim Biophys Acta 1852(7):1298–310.

Yan Y, Qiao S, Kikuchi C, Zaja I, Logan S, Jiang C, Arzua T, Bai X. 2017. Propofol Induces Apoptosis of Neurons but Not Astrocytes, Oligodendrocytes, or Neural Stem Cells in the Neonatal Mouse Hippocampus. Brain Sci 7(10).

Yu D, Li L, Yuan W. 2017a. Neonatal anesthetic neurotoxicity: Insight into the molecular mechanisms of long-term neurocognitive deficits. Biomed Pharmacother 87:196–199.

Yu S, Patchev AV, Wu Y, Lu J, Holsboer F, Zhang JZ, Sousa N, Almeida OF. 2010. Depletion of the neural precursor cell pool by glucocorticoids. Ann Neurol 67(1):21–30.

Yu S, Zutshi I, Stoffel R, Zhang J, Ventura-Silva AP, Sousa N, Costa pS, Holsboer F, Patchev A, Almeida OF. 2017b. Antidepressant responsiveness in adulthood is permanently impaired after neonatal destruction of the neurogenic pool. Transl Psychiatry 7(1):e990.

Zhao C, Deng W, Gage FH. 2008. Mechanisms and functional implications of adult neurogenesis. Cell 132(4):645–60.

Zhu C, Gao J, Karlsson N, Li Q, Zhang Y, Huang Z, Li H, Kuhn HG, Blomgren K. 2010. Isoflurane anesthesia induced persistent, progressive memory impairment, caused a loss of neural stem cells, and reduced neurogenesis in young, but not adult, rodents. J Cereb Blood Flow Metab 30(5):1017–30.

